# Combinatorial Immunotherapies Overcome MYC-Driven Immune Evasion

**DOI:** 10.1101/2021.05.07.442684

**Authors:** Joyce V. Lee, Filomena Housley, Christina Yau, Daniel Van de Mark, Rachel Nakagawa, Golzar Hemmati, Grace A. Hernandez, Juliane Winkler, Yibing Zhang, Susan Samson, Carole Baas, Laura J. Esserman, Laura J. Van ‘T Veer, Hope S. Rugo, Mehrdad Matloubian, Andrei Goga

## Abstract

For many human cancers, including triple negative breast cancer (TNBC), a modest number of patients benefit from immune checkpoint inhibitors, and few experience cancer remission^1^. Expression of programed death-ligand 1 (PD-L1), tumor immune infiltration, or tumor mutation burden have been widely investigated for predicting cancer immunotherapy response^1-5^. Whether specific oncogenes diminish response to immunotherapy^6-10^ and whether these effects are reversible remains poorly understood. We predicted that *MYC*, an oncogene that is frequently overexpressed^11,12^ and is associated with worse prognosis^12^, may predict immunotherapy response in patients with TNBC. Here, we report that MYC-elevated TNBCs are resistant to immune checkpoint inhibitors. Using mouse models of TNBC and patient data we report that MYC signaling is associated with low tumor cell PD-L1, low overall immune cell infiltration, and low tumor cell MHC-I expression. Restoring interferon signaling in the tumor reduces MYC expression and increases MHC-I expression. By combining a TLR9 agonist and an agonistic antibody against OX40 with anti-PD-L1, most mice experience complete tumor regression and are protected from new TNBC tumor outgrowth. Our findings demonstrate that MYC-dependent immune evasion is reversible and druggable, and if strategically targeted, may improve outcomes for patients treated with immune checkpoint inhibitors.

---

To understand how MYC might alter response to immune checkpoint inhibitors, we initiated our studies using a genetically engineered mouse model of triple negative breast cancer (MTB/TOM)^13^, where MYC expression in breast epithelial cells can be switched on and off with doxycycline. Tumors that arose from this model can be propagated via transplantation into the fourth mammary fat pad of wild-type FVBN mice. MYC expression was detected in the tumors while mice were fed doxycycline (MYC-ON) and a loss of MYC in the tumor within 3 days of taking mice off doxycycline chow (MYC-OFF) (Extended Data Fig.1a).

A prior study found that MYC upregulates tumor cell PD-L1, a cell surface molecule that dampens the adaptive immune response, in a mouse model of MYC-driven lymphoma^14^, suggesting blocking PD-L1 therapy might be effective in MYC-driven cancers; however, in a KRas^*G12D*^/MYC-driven lung cancer mouse model, anti-PD-L1 was ineffective^10^, suggesting that PD-L1 expression may be cancer type specific. These conflicting results prompted us to investigate the role of PD-L1 in MYC-driven TNBC.

We allowed each animal’s tumor to grow to 10 mm in length and began anti-PD-L1 treatment (Extended Data Fig. 1b). As a single agent, anti-PD-L1 failed to slow tumor growth (Fig. 1a). To characterize PD-L1 expression in the tumors, we dissociated tumors and used flow cytometry. While cell surface PD-L1 expression was observed on the tumor myeloid cells (CD45+, monocytes and dendritic cells), it was absent on the tumor cells (CD45-, EPCAM+) (Fig.1b).

**Fig. 1.**
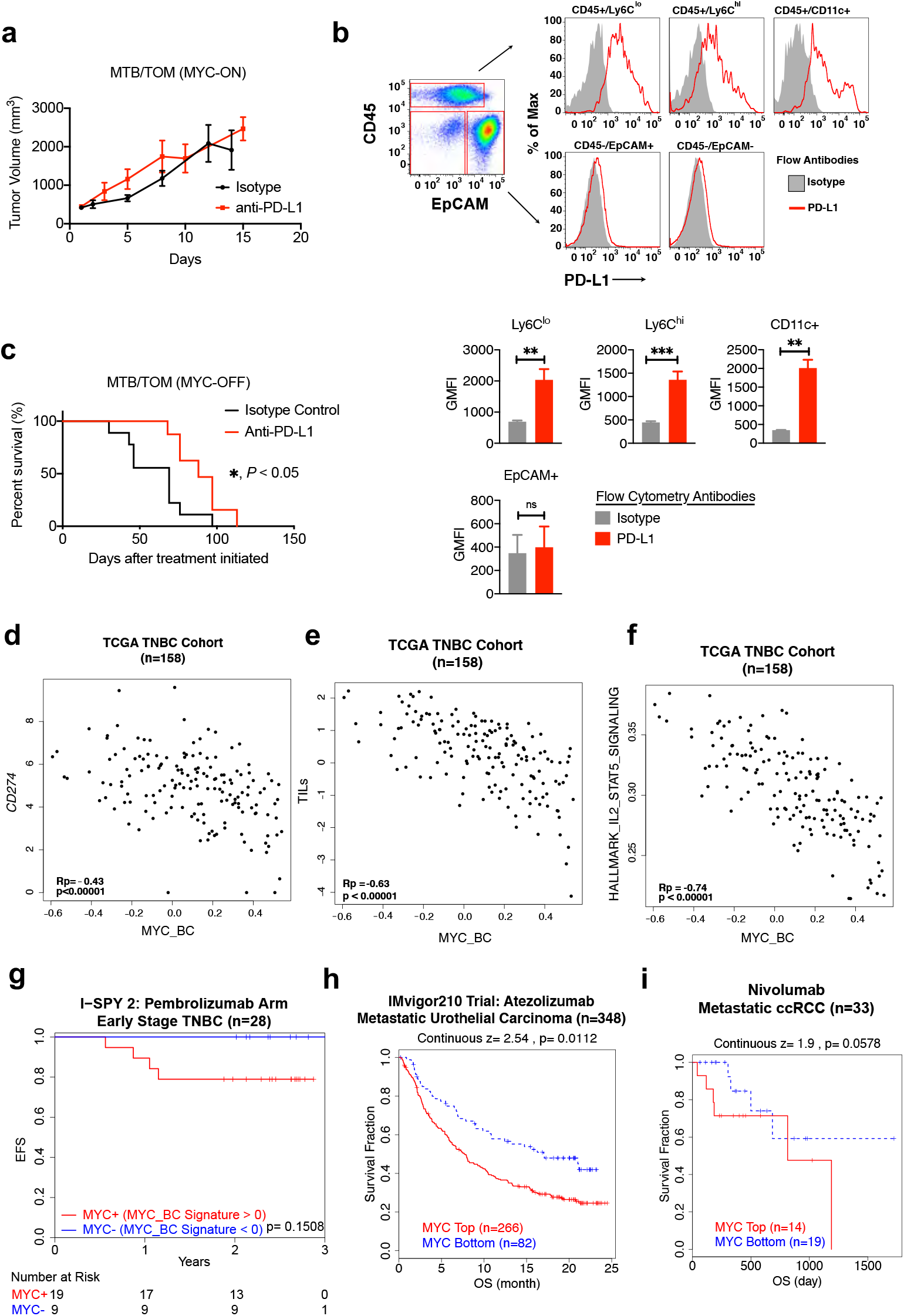
MYC predicts poor response to immune checkpoint inhibitors. **(a)** Average tumor volume over time for MTB/TOM tumors treated with anti-PD-L1 or isotype antibody control while fed doxycycline chow (MYC-ON state). n=3-4 tumors per group, mean ± S.E.M. **(b)** Top: Representative flow cytometry plots comparing mouse PD-L1 protein expression in CD45- or CD45+ populations in MTB/TOM tumors. On the right, PD-L1 expression (red) and isotype antibody (gray) displayed as percent of maximum (modal). Bottom: Bar graphs displaying average geometric mean fluorescence intensity (GMFI) for PD-L1 in each population. n=3-6 tumors, Mean ± S.E.M., Unpaired t-test. **, p < 0.01; ***, p < 0.001; ns, p ≥ 0.05. **(c)** Survival analysis for time to recurrence in MTB/TOM tumors off doxycycline. Median time to ethical study endpoint was 69 days in MYC-OFF + isotype antibody (n=9) and 88 days in MYC-OFF + anti-PDL1 (n=8). Log rank test; p = 0.017. (d-f) Scatterplots of **(d)** *CD274* expression, **(e)** the tumor infiltrating leukocyte (TIL), and (**f)** IL2/STAT5 Hallmark gene expression signatures against the MYC_BC signature in the TCGA TNBC cohort (n=158). Pearson correlation (Rp) and p values are shown (bottom left). **(g)** Kaplan-Meier curves of event-free survival for 28 patients with HER2 negative and hormone receptor (HR) negative tumors treated on the Pembrolizumab arm of the I-SPY 2 TRIAL dichotomized by MYC_BC signature. (**h-i**) Survival analysis using *MYC* expression from pre-treatment tumors pulled out of the TIDE database: **(h)** Kaplan-Meier curves of overall survival in 384 patients with metastatic urothelial carcinoma from Mariathasan et al., 2018. **(i)** Kapaln-Meier curves of overall survival in 33 patients with metastatic clear cell renal cell carcinoma (ccRCC) from Miao et al., 2018.

We tested if dampening MYC expression could improve response to anti-PD-L1 therapy. Once tumors grew to 10 mm in length, we switched MYC off in all tumors by removing doxycycline from the mice’s diet and concurrently started them on anti-PD-L1 or isotype control antibody (Extended Data Fig. 1c); the animals remained off doxycycline through the end of the study. Tumors shrank initially, but all eventually recurred (Extended Data Fig. 1d). Nevertheless, by combining MYC inactivation with anti-PD-L1 therapy, we significantly delayed tumor recurrence, extending median survival by 25% more than isotype treated (Fig. 1c). Taken together, MYC-driven breast tumors do not express PD-L1 and do not respond to anti-PD-L1 alone, but inactivation of MYC in combination with anti-PD-L1 delayed tumor recurrence and extended survival in mice.

To determine if MYC signaling is correlated with specific immune cell types, which might impact response to anti-PD-L1, we turned to tumor samples from patients with TNBC from the Cancer Genome Atlas (TCGA) dataset. To model gene expression in the breast tumor microenvironment, we derived a MYC signature specific to breast cancer using published gene expression data from multiple MYC-driven mouse models of breast cancer. First, we selected the significantly up- and down-regulated genes in the MYC expression subtype (MYC^ex^) derived from the TgMMTV-Myc mouse and TgWAP-Myc mouse from Pfefferle and colleagues^15^, and the genes that were significantly altered by MYC in MTB/TOM model published by our lab^16^ (Extended Data Fig. 2a). We identified 530 shared mouse genes regulated by MYC in breast cancer. These mouse genes were then matched to their corresponding human gene IDs to generate an *in vivo* MYC-driven breast cancer (MYC_BC) signature. We confirmed that the MYC_BC signature is highly correlated with a previously published *in vitro*-derived MYC signature^12,17^ (Extended Data Fig. 2b) and the Molecular Signatures Database (MSigDB) Hallmark MYC Targets signature (Extended Data Fig. 2c).

Patients with a high MYC_BC signature had less *CD274* expression (Fig. 1d). Next, we explored how the MYC_BC signature correlated with published immune cell signaling signatures in the TCGA TNBC patient cohort. Overall, tumors with a high MYC_BC signature were associated with reduced tumor infiltrating leukocytes (TILs) (Fig. 1e). The MYC_BC signature was negatively correlated with published immune cell type signatures^18^, showing fewer T cells, B cells, macrophages, and NK cells with MYC activation (Extended Data Fig. 3). The MYC signature also associated with low T-cell cytokine signaling (IL2/STAT5) (Fig. 1f).

## MYC associates with poor survival after immune checkpoint inhibition

Given the low immune signatures associated with MYC-elevated human tumors, we predicted that for patients on immune checkpoint inhibitors, those with a high MYC signature score would associate with worse outcomes. We looked in a small cohort of patients with early stage TNBC treated with pembrolizumab, a monoclonal antibody therapy targeting PD-1, plus standard chemotherapy in the neoadjuvant setting from the I-SPY 2 TRIAL^19^. At the median follow up time of 2.4 years, all observed recurrences were among the patients with a high MYC_BC signature, and no recurrence was seen in those patients with a low MYC_BC signature (Fig. 1g). Though the study was underpowered, the clear trend pointed us to MYC as a potential predictive biomarker to evaluate outcomes of patients receiving immune checkpoint inhibitors. This prompted us to look at other published immunotherapy datasets though the Tumor Immune Dysfunction and Exclusion (TIDE) platform^20,21^. Because the database does not have trials involving breast cancer, we queried for *MYC* expression rather than using the MYC_BC signature. Most trials had small numbers of patients, nevertheless we profiled urothelial cancer^22^ and renal cell carcinoma (ccRCC)^23^, tumor types with MYC amplification^24,25^ or overexpression^24,26^. In urothelial cancer, patients had shorter median overall survival after anti-PD-L1 if their pre-treatment tumors expressed high *MYC* (p = 0.0112) (Fig. 1h). A trend toward diminished overall survival was also found with elevated *MYC* in patients with ccRCC (p = 0.0578) (Fig. 1i).

## MHC-I is downregulated in MYC-activated tumors

Though PD-L1 was present in tumor associated immune cells (Fig 1b), the poor tumor response to anti-PD-L1 (Fig 1a) led us to postulated that other factors might explain the observed immune evasion. This prompted us to explore the cellular programs that might contribute to poor response in the MYC-ON state. Tumor-specific antigen presentation by MHC class 1 (MHC-I) is necessary for T-cell recognition and cytotoxicity^27^. Somatic loss of heterozygosity in antigen presentation machinery is associated with resistance to immunotherapy in patients with melanoma^28^ and lung cancer^29,30^. Genetic deletion of a key MHC-I component, β2-microglobulin (*B2m*), abrogates the therapeutic effect of anti-PD-1 in mouse melanoma^31^.

We decided to explore the expression of *B2M* and other antigen presentation genes in the context of MYC and immunotherapy. Although a link between MYC family proteins and MHC-I heavy chain gene expression was described over 30 years ago in neuroblastoma and melanoma cell lines^32,33^, it has not been shown in breast cancer, and its significance for immune checkpoint inhibitor response has not been investigated in human patient samples or *in vivo* models.

Patients with a high MYC_BC signature had lower expression of genes important for MHC-I expression (Extended Data Fig. 4), such as *B2M* (Fig. 2a) and *NLRC5* (Fig. 2b), the major transactivator of MHC-I genes^34,35^. We further validated that MYC is accompanied by a loss of MHC-I gene expression in human cells by examining gene expression in an immortalized, non-tumorigenic human mammary epithelial cell line, MCF10A. MCF10A cells overexpressing ectopic MYC^36^ displayed lower expression of antigen presentation machinery genes than parental MCF10A cells, indicating that MYC overexpression is sufficient to drive the repression of MHC-I related genes in human breast epithelial cells (Extended Data Fig. 5).

**Fig. 2.**
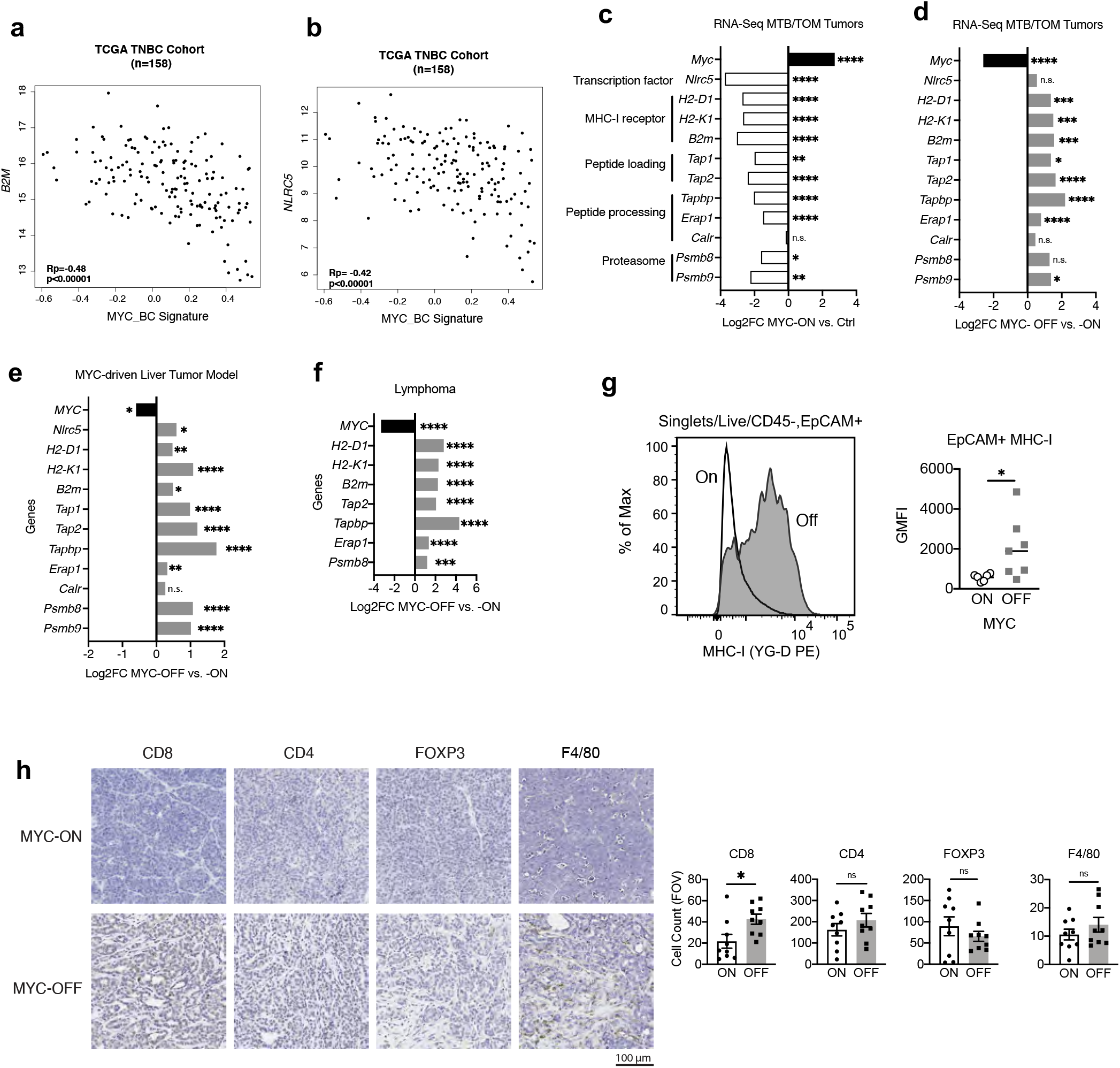
MYC suppression of MHC-I expression is reversible. **(a-b)** Scatter plot showing a negative correlation (Rp, Pearson’s correlation coefficient) between the expression of **(a)** *B2M* gene and the MYC signature in TNBC patients n=158 or **(b)** *NLRC5* gene and the MYC signature in TNBC patients n=158. **(c)** RNA-seq data in MTB/TOM tumors (MYC-ON) compared to normal mammary tissue (Ctrl). Adjusted p values reported. **(d)** RNA-seq in MTB/TOM tumors 72 hours off-doxycycline (MYC-OFF) compared to animals on doxycycline (MYC-ON). Adjusted p values reported. **(e)** Representative histogram for MHC-I expression by flow cytometry in MYC-ON and MYC-OFF states. Adjacent bar graph displaying geometric mean fluorescence intensity (GMFI) for MHC-I in MYC-ON (n=6) and MYC-OFF (n=7) states. Line represents median, data points represent individual animals, Mann-Whitney test. **(f)** Immunohistochemistry staining of immune cell markers within tumor sections in MYC-ON and MYC-OFF state. Images displayed are at 20X. Adjacent bar graphs displaying cumulative counts per field of view (FOV). n=3 animals per group, 3 FOV analyzed per tumor, mean ± S.E.M., Mann-Whitney test.

Next, we compared the expression of antigen presentation genes in the MYC-ON mouse breast tumors to normal mouse mammary glands (Ctrl). The presence of MYC significantly downregulated multiple genes important for antigen presentation by MHC-I (Fig. 2c). Remarkably, most antigen presentation genes were re-expressed within 3 days of inactivating MYC (MYC-OFF) (Fig. 2d), suggesting that the expression of these genes is dependent on MYC. We investigated whether MYC exhibited similar effects on MHC-I genes in a published liver cancer model^37^ and lymphoma model^16^; indeed, turning MYC off in these models also resulted in an increase in antigen presentation genes (Fig. 2e, f), suggesting that the downregulation of antigen presentation genes is dependent on MYC in multiple conditional transgenic cancer models.

To validate the reduction of MHC-I gene expression by MYC, we investigated the cell surface expression of MHC-I in MTB/TOM tumors after MYC inactivation. Tumor cells (CD45-, EPCAM+) in the MYC-OFF state displayed significantly more surface MHC-I protein than MYC-ON tumor cells (Fig. 2g). We also observed increased tumor infiltration of CD8+ T cells in the MYC-OFF state, while the infiltration by other immune subtypes was not significantly altered (Fig. 2h). Together, these data demonstrate MYC-ON tumors have low MHC-I expression and turning MYC off rescued MHC-I cell surface expression and concurrently increased CD8+ T-cell tumor infiltration. These two phenotypes observed upon MYC-inactivation provide a possible explanation as to why turning MYC off improved response to anti-PD-L1 therapy in Figure 1. These studies demonstrate that the transcriptional regulation and cell surface expression of MHC-I is reversible and thus, targetable.

### Interferons rescue MHC-I in MYC-elevated tumors

Without approved drugs that directly inhibit MYC, we dug deeper into the MTB/TOM RNA-seq data for clues toward targeting MYC-driven breast cancers. We assembled a list of genes downregulated by MYC *in vivo* to discover new targetable pathways. We queried genes downregulated in the MYC-ON state and upregulated in MYC-OFF state. This yielded 1328 genes interrogated using MSigDB^38^ (Extended Data Fig. 6). MYC-downregulated pathways were enriched for immune processes (Extended Data Table 1), indicating that restoring immune infiltration could improve efficacy of anti-PD-L1 therapy. The interferon response pathways were among the most suppressed pathways in MYC-activated tumors (Extended Data Table 1). We examined the hallmark interferon-alpha and interferon-gamma pathways in the TCGA TNBC tumors; we found higher MYC signature in tumors with less interferon signaling (Fig. 3a, b).

**Fig. 3.**
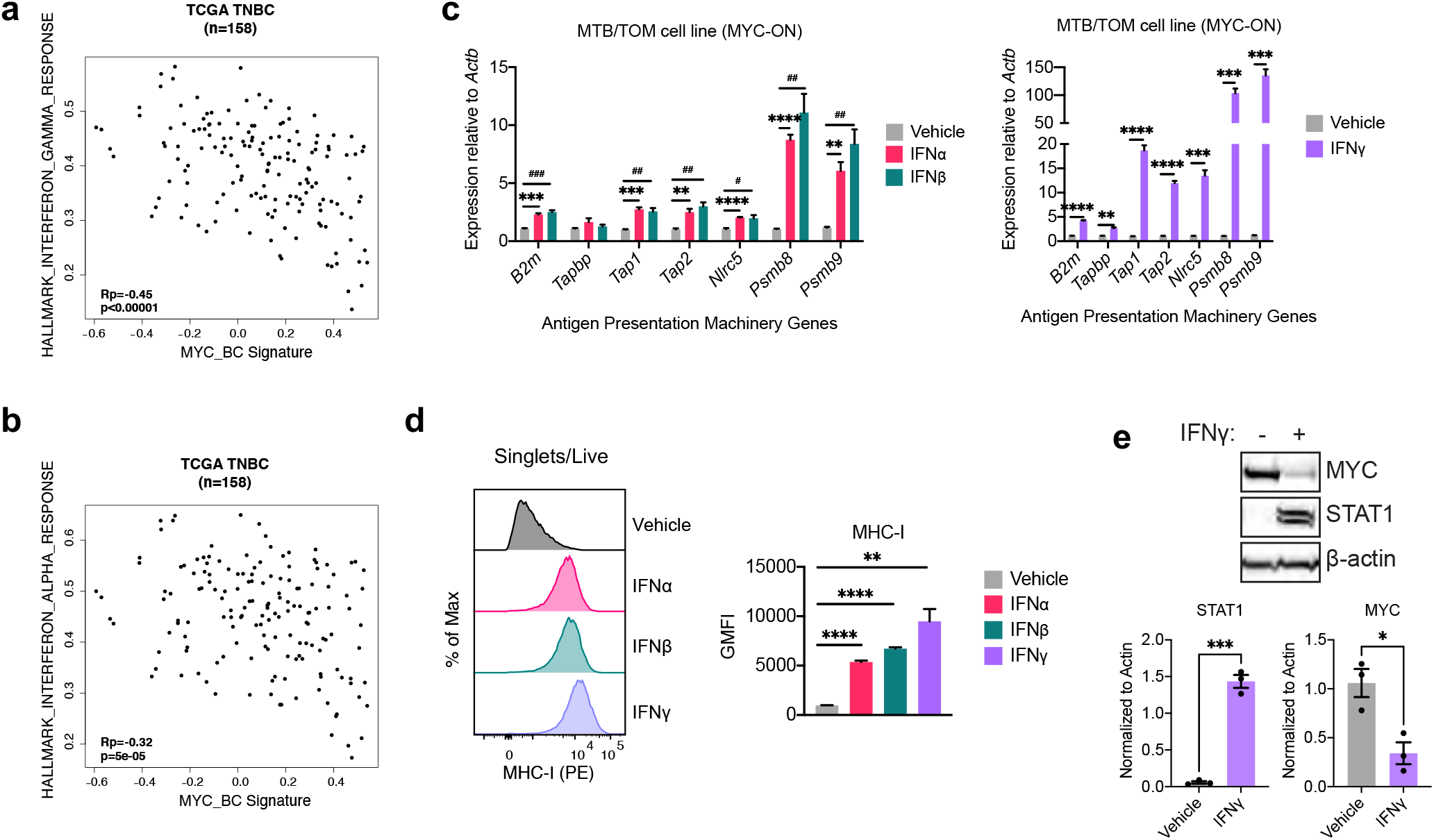
Interferon signaling rescues MYC suppression of MHC-I. **(a-b)** Scatter plot showing a negative correlation between the expression of **(a)** Hallmark Interferon Gamma Response signature and the MYC signature in TNBC patients n=158 or **(b)** Hallmark Interferon Alpha Response signature and the MYC signature in TNBC patients n=158. Pearson correlation (Rp) and p values are shown (bottom left). **(c)** Gene expression analysis after 72 hours of vehicle, interferon alpha, beta (left) or gamma (right) treatment in the presence of doxycycline, in MTB/TOM cells grown *in vitro* culture. Representative experiment with 3 samples per condition shown. Mean± S.E.M, unpaired t-test with Benjamini-Hochberg FDR=0.05. # or *, p < 0.05; ## or **, p < 0.01; +++, ###, or ***, p < 0.001; ****, p < 0.0001. Experiment repeated with 3 independent cell passage replicates. **(d)** Representative histogram of cell surface expression of MHC-I as detected by flow cytometry after 72 hours of treatment with interferons in MTB/TOM cell culture, in the presence of doxycycline. Adjacent bar graph displaying geometric mean fluorescence intensity (GMFI) for MHC-I. Mean± S.E.M, unpaired t-test, data points represent 3 independent samples. **(e)** Representative western blot showing MYC levels after 72 hours of treatment in the MTB/TOM cell line. Quantification of results from 3 independent cell passage experiments displayed below. Unpaired t-test. *, p < 0.05; ***, p < 0.001.

Canonically, MHC-I is expressed robustly upon interferon signaling^39^, but whether this is true in breast cancer is unknown. We tested whether MYC-high tumors could still repress MHC-I after exogenous interferon stimulation. We exposed a cell line derived from MTB/TOM tumors to type I interferons (IFNα or IFNβ) or type II interferon (IFNγ), while in the MYC-ON state. Following exposure, we observed the induction of several antigen presentation genes, with IFNγ inducing a greater change than the type I interferons (Fig. 3c), and a dramatic increase of MHC-I protein on the cell surfaces of the MTB/TOM tumor cells (Fig. 3d). We tested whether the re-expression of MHC-I was due to changes in MYC. Even when the MTB/TOM cells were maintained on doxycyline, we detected a decrease in total levels of MYC protein following IFNγ by western blot (Fig. 3e). These studies suggest that restoring interferons is one way to overcome MYC suppression of MHC-I protein expression on tumor cells.

### CpG/aOX40 improves anti-PD-L1

Activation of pattern recognition receptor pathways, such as toll-like receptor 9 (TLR9), leads to the local production of interferons in the tumor microenvironment. A synthetic TLR9 agonist, unmethylated CpG oligodeoxynucleotides (CpG-ODN or CpG), can stimulate plasmacytoid dendritic cells (pDC) to produce IFNα and IFNβ in the local microenvironment, activating B cells and attracting natural killer (NK) cells; this cascade upregulates production of IFNγ and subsequently can attract antigen-specific CD8+ T-cells^40^.

We asked whether a concerted interferon response induced by CpG would increase MHC-I presentation in the MYC-driven mouse mammary tumors. After a single intratumoral administration of CpG directly into MYC-ON tumors, we observed more MHC-I expression on the tumor cells (Fig. 4a, Extended Data Fig. 7).

**Fig. 4.**
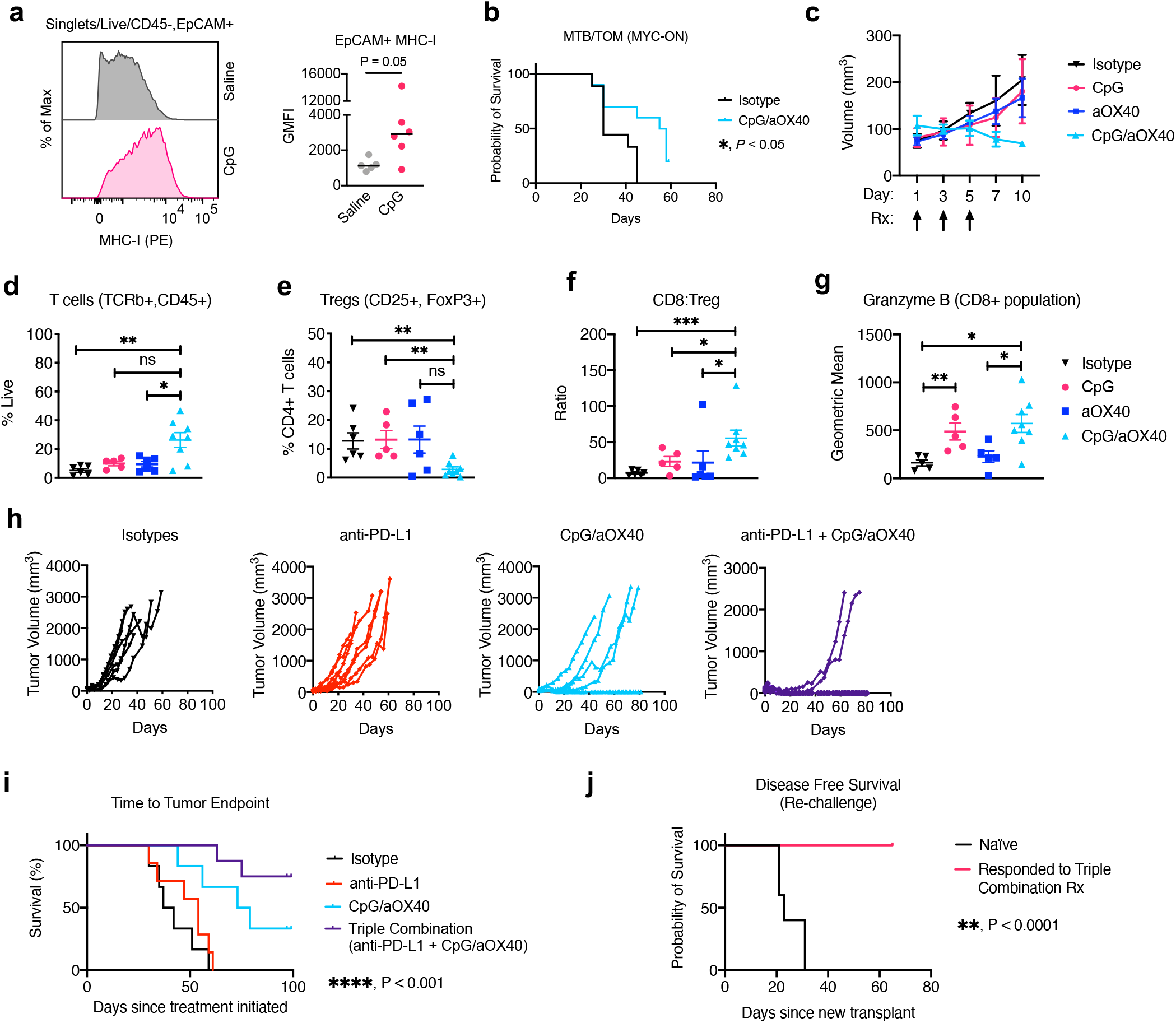
CpG/anti-OX40 enhances anti-PD-L1 *in vivo*. **(a)** Representative histogram of cell surface expression of *in vivo* MHC-I detected by flow cytometry, 48 hours after CpG injection or saline injection directly into tumors 5 mm in length. Adjacent bar graph displaying geometric mean fluorescence intensity (GMFI). n=5-6 animals per treatment group. Line represents median, data points represent individual animals, Mann-Whitney test. **(b)** Survival analysis for MTB/TOM treated with CpG/anti-OX40 (aOX40). Median time to endpoint was 56.5 days for CpG/aOX40 group (n=10) and 30 days for the isotype vehicle control group (n=9). Log rank test; p= 0.0154. **(c)** Tumor volume in first 10 days, following treatments on day 1, 3, and 5, indicated by arrows. **(d-g)** Flow cytometry analysis of immune cells found in the tumor at day 10. n=6 isotype, n=5 CpG, n=6 anti-OX40, n=8 CpG/aOX40. **(d)** Percent of live cells in the tumor that are immune cells. **(e)** Percent of CD4+ (gating: Singlets/Live/TCRβ+, CD45+) cells that are CD25+ and FOXP3+. **(f)** Ratio of CD8+ (gating: Singlets/Live/TCRβ+, CD45+, CD8+) counts to Treg counts (gating: Singlets/Live/TCRβ+, CD45+, CD25+, FOXP3+). **(g)** Geometric mean fluorescence intensity for intracellular Granzyme B in the CD8+ population (gating: Singlets/Live/TCRβ+, CD45+, CD8+). Mann-Whitney test. For all panels: Mean± S.E.M; *, p < 0.05; **, p < 0.01; ***, p < 0.001; ****, p < 0.0001. **(h)** Tumor volumes since the start of treatment: isotype (n=6), anti-PD-L1 (n=7), CpG/aOX40 (n=6), and triple combination (n=9). **(i)** Survival curves for animals in panel h. (**j)** Disease-free survival (defined as time to palpable tumor) in mice transplanted with a new MTB/TOM tumor on the left side, fourth mammary fat pad. n=7 in control group and n=5 in responded to therapy group. For h-i: log rank test; **, p < 0.01; ****, p < 0.0001.

Intratumoral CpG combined with systemic anti-PD-L1 did not extend survival beyond CpG alone (Extended Data Fig. 8), suggesting another form of immunosuppression is involved. A recent study revealed that CpG induces the expression of OX40, a co-stimulatory molecule, on CD4+ T-cells (including suppressive T-regulatory cells, Tregs)^41^. Stimulation of Tregs through OX40 impairs their function, which is critical for tumor shrinkage in the spontaneous polyomavirus middle T-antigen breast cancer mouse model^41^. We decided to test whether a combination of CpG and agonistic antibody against OX40 (aOX40), could delay tumor progression in MTB/TOM tumors. We administered intratumoral injections of CpG oligo and anti-OX40 (CpG/aOX40) in MYC-ON tumors (5 mm) every other day for a total of 3 injections and then monitored tumors to study endpoint. CpG plus anti-OX40 treatment delayed tumor progression and increased median survival by 86% (Fig. 4b).

To discern the effects of CpG/aOX40 on the tumor microenvironment, we examined the immune composition at day 10, several days after administering the last dose. At this time point, the anti-tumor effects of CpG/aOX40 emerged (Fig. 4c). We detected a dramatic increase in immune infiltration as detected by immunohistochemistry staining (Extended Data Fig. 9), including total T-cells (CD3+), specifically, CD4+ and CD8+ T-cells, and a trend toward more macrophages (F4/80) (Extended Data Fig. 9). These results were confirmed and further characterized by flow cytometry. We found more T-cells (CD45+, TCRβ+) in the tumors given CpG/aOX40 compared to tumors given isotype control or aOX40 alone (Fig. 4d). Concurrently, we observed a lower proportion of Tregs, (CD25+, FoxP3+), within the CD4+ T-cell population, for CpG/aOX40 treated tumors (Fig. 4e). Compared to single agent treated tumors, CpG/aOX40 treated tumors displayed a higher CD8+ T-cell to Treg ratio (Fig. 4f). Furthermore, the CD8+ T-cells (CD45+, TCRβ+/CD8+) expressed greater amounts of granzyme B molecules in the CpG group and CpG/aOX40 group, compared to the isotype treated tumors (Fig.4g), suggesting that the specific treatments increased CD8 T cell functionality and ability to initiate tumor killing. Overall, CpG/aOX40 treatment sustained high CD8+ T cell:Treg and granzyme B production.

Given both the quantitative and qualitative changes in the intratumoral CD8+ T cells and increased tumor MHC-I expression following CpG/aOX40 treatment in MYC-ON tumors, we reasoned that this approach would improve the sensitivity to anti-PD-L1 therapy. We therefore tested CpG/aOX40 together with anti-PD-L1. Once tumors reached 5 mm, animals were randomized into the 4 treatment arms (Extended Data Fig. 10). Isotype antibody treated and anti-PD-L1 alone treated animals displayed rapid tumor progression (Fig. 4h). In the CpG/aOX40 group, one third of the animals had tumors regressed (Fig 4h, 4i). Remarkably, addition of anti-PDL1 (triple combination) resulted in complete and long-term regression of tumors in 75% of the animals, at 100 days post-treatment initiation (Fig. 4h, 4i).

We challenged the mice that fully regressed tumors with new tumors transplants in their contralateral mammary fat pads. FVBN mice that had not received prior tumor transplantation nor previous therapy served as controls for tumor growth. All animals that responded to the initial therapy successfully eliminated growth of new tumor transplants (Fig. 4j), demonstrating that the combination therapy led to a durable immune response that protected mice from establishing new tumor outgrowth.

## Discussion

Immune checkpoint inhibitors represent novel and promising treatments for patients. Efficacy is seen in patients with PD-L1 positive tumors, but few patients achieve remission^1^. We observed that anti-PD-L1 was ineffective in MYC-driven breast cancer models, even though PD-L1 was expressed on tumor-associated myeloid cells. This highlighted that in the context of MYC, additional tumor immune evasion mechanisms were relevant.

The low efficacy of anti-PD-L1 in MYC-elevated cancers is partly due to suppression of MHC-I genes and low tumor immune cell infiltration in patient tumors and mouse models. The MYC-dependent suppression of T-cell infiltration seen here is supported by previous findings in a KRas^*G12D*^/MYC-driven lung cancer model and pancreatic cancer model^10,42^. We further demonstrate that inactivation of MYC in the MTB/TOM TNBC mouse model restored MHC-I cell surface expression, CD8+ T cell infiltration, and response to anti-PD-L1. Collectively, our data emphasizes that MYC-dependent deregulation of MHC-I and immune exclusion is reversible and thus, druggable by increasing interferon signaling. In lung cancer cells, EZH2 inhibitor^43^ and HDAC inhibitor^44^ have been demonstrated to restore MHC-I *in vitro*, suggesting these drugs might also increase MHC-I in lung tumors. Here, we demonstrate *in vivo* that administration of a single, low dose of CpG alone was sufficient to bypass MYC-dependent repression of breast tumor cell MHC-I, and it increased the fraction of CD8+ T-cells that can induce tumor cell apoptosis (production of GrB). Importantly, a short interval treatment of CpG/aOX40 significantly reduced the fraction of immunosuppressive Tregs within mouse MYC-driven TNBC tumors and maintained cytotoxic CD8+ T cell infiltration. When combined with anti-PD-L1, a majority of tumors fully regressed, suggesting that local immunostimulation is effective in MYC-driven tumors.

Our study suggests that patients with MYC-elevated tumors will likely require additional therapies to achieve optimal survival benefit with immune checkpoint inhibitors. This is highlighted in our analysis of event free survival (EFS) for patients with TNBC in the pembrolizumab arm of the ISPY-2 TRIAL^19^, which revealed that patients with a high MYC signature in their pre-treatment tumors experienced tumor relapse or metastasis sooner than patients with a low MYC signature. All patients received chemotherapy together with immune checkpoint blockade, suggesting that chemotherapy is not sufficient to cause an immunogenic response in tumors with high MYC-signaling. MYC may also be an important predictor of outcomes for immune checkpoint inhibitor therapy in other tumor types (such as urothelial bladder cancer and ccRCC). Clinical trials like the Phase II I-SPY2 and the Phase III ILLUMINATE-301 testing immune checkpoint inhibitors and TLR9 agonists are ongoing and would likely benefit patients with MYC-elevated tumors. We believe the implementation of a MYC_BC signature will be critical to identifying patients that are at high risk for progression on immune checkpoint inhibitors, and a focus on therapeutics that rescues MHC-I expression and enhances anti-tumor immune cell infiltration will specifically improve the efficacy of immune checkpoint inhibitor therapies like anti-PD-1 and anti-PD-L1 for these patients.

## Methods

### Tumor initiation in mice

Viably frozen MTB/TOM tumors (lab lines: A and B), generated from MMTV-rtTA/TetO-MYC mice on the FVBN background, were divided into fragments (∼8 mm^3^) and one piece was implanted into the cleared 4^th^ mammary fat pad of each 4-week-old female FVBN mice (Jackson Laboratory Stock 001800). Mice were maintained on doxycycline diet (Bio-Serv #S3888) starting 1 day before transplant surgery. All mice were maintained at UCSF rodent barrier facilities. All procedures were approved by UCSF Institutional Animal Care and Use Committee (IACUC) under protocol number AN184330-01.

### Tumor studies

For PD-L1 monotherapy studies, animals were randomized to treatment groups when tumors reached 10 mm; for animals off doxycycline (MYC-OFF), animals were moved into new cages with provided standard chow. For combinatorial therapy experiments, animals were randomized to treatment arms once tumors reached 5 mm in length. Saline was used to dilute the drugs. The drugs for each arm are: CpG (IDT) was delivered intratumorally with an insulin syringe at 50 µg in 0.05 mL per injection; anti-OX40 (Bio X Cell #BE0031) was delivered intratumorally with an insulin syringe at 8 µg in 0.05 mL; CpG/anti-OX40 together was delivered in a single injection containing 50 µg of CpG and 8 µg of anti-OX40 in 0.05 mL. Anti-PD-L1 (Bio X Cell #BE0101) was delivered by I.P. at 0.2 mg in 0.2 mL per injection. For experiments involving anti-OX40 and/or anti-PD-L1, the control group was treated with the respective isotype antibodies (Bio X cell #BE0290, #BE0090). For experiments involving CpG, the control group was given saline. Ethical endpoint was defined as tumor length of 20 mm in any direction. Tumor volume was calculated as (length) ÷ 2 × (width)^2^.

### Flow cytometry

Each tumor was minced with a clean blade and then dissociated in 5 mL RPMI (Gibco) containing 1 mg/mL collagenase IV (Gibco), 0.1 mg/mL DNase (Roche), 2% heat inactivated FBS (Gibco), and 10 mM HEPES at 37 °C with constant agitation of 180 rpm for 25 minutes. Digested tissue was diluted with 30 mL of cold PBS and poured through a 70-micron nylon mesh strainer (Fisher) for tumor cell analyses or 40-micron nylon mesh strainer for immunophenotyping. Cells were pelleted at 220-300 x g and resuspended in 5 mL RBC lysis buffer (BioLegend) at room temperature. After 5 minutes, the cells were diluted with 25 mL of FACS buffer (HBSS with 10mM EDTA and 2% heat inactivated FBS), pelleted, and resuspended in < 3 mL of PBS for cell counts. Cells were stained with the antibody panel (see supplementary table) for 30 minutes on ice, covered. Cells were washed with PBS and stained with fixable Near-IR live/dead stain (Molecular Probes) at 1:1,000 for 15 minutes at room temperature. For FOXP3 and Granzyme B staining, cells were fixed and permeated with a transcription factor staining buffer set (Invitrogen 00-5523-00) following staining for extracellular proteins. Cells were washed and resuspended in FACS buffer for analysis on a BD Dual Fortessa and analyzed with FlowJo 10. For MTB/TOM cells grown in culture, the same reagents and protocols were used after cells were lifted off the plate with a cell lifter (Corning), but the RBC lysis step was omitted. All experiments were compensated with single color controls and gating was determined by full panel minus one antibody or isotype antibody controls. All antibodies, unless otherwise indicated, were used at 1:100. A list of antibodies is in Supplementary Table 1.

### Histology and immunohistochemistry (IHC)

Tissues were fixed in paraformaldehyde (Electron Microscopy Sciences, #15700), diluted to 4% with PBS, for 16-20 hours and then moved to 70% ethanol. Samples were further processed by HistoWiz Inc, using their standard operating procedure and automated immunohistochemistry staining workflow with their in-house validated antibody list (published June 2018).

### Immunohistochemistry quantification

For each tumor and stain, three representative fields taken at 40X were imported for analysis using Fiji/ImageJ tool for color deconvolution for H DAB. The brown channel (colour-2) was further selected for analysis. The threshold was standardized for each staining and set for all tumor samples. For T cell specific stains (CD3, CD4, CD8, FOXP3), H DAB positive particle counts were recorded. For F4/80 specific staining, the percentage of H DAB positive area (output % Area) was recorded.

### Western blot

Cells were grown in 6-well dishes and grown to 90% confluence on the day of harvest. Cells were quickly washed once with cold PBS and lysed with Laemmli buffer (60 mM Tris-HCl, pH 6.8, 1 μM DTT, 2% w/v SDS) supplemented with protease inhibitor cocktail (Roche) and phosSTOP (Roche) or RIPA buffer (Thermo) supplemented with protease inhibitor cocktail and phosSTOP. Protein extracts were quantified with DC Protein Assay (Bio-Rad) and prepared with NuPage sample loading buffer and reducing agent (Invitrogen). Proteins were resolved with the Bolt 4-12% Bis-Tris gel and buffer system (Invitrogen) and transferred onto nitrocellulose membranes using iBlot2 program P0 (Invitrogen). Membranes were blocked with 5-10% non-fat milk (Rockland) in TBST and incubated with primary antibodies diluted in 5% non-fat milk TBST overnight on a 4°C shaker. Membranes were washed with TBST and incubated with horseradish peroxidase (HRP)-conjugated secondary antibodies in TBST the next day. Signals were captured with ECL Prime (GE) or Visualizer (Millipore) on a Bio-Rad GelDoc system and Image Lab software. Images were exported to ImageJ for quantification using ‘Analyze > Gels’. The following antibodies were used: Anti-ß-actin (1:10,000, sc-47778 HRP, Santa Cruz Biotechnology), anti-c-MYC (1:1,000, clone Y69, ab32072, Abcam), anti-STAT1 (1:1,000, #9172, Cell Signaling), Anti-Rabbit IgG (1:10,000, #7074, Cell Signaling).

### RT-qPCR

Cells were seeded on 6-well dishes and grown to 90% confluence on the day of harvest. Cells were quickly washed once in cold PBS, and total RNA was isolated using Trizol (Invitrogen) per manufacturer’s instructions. 2 μg of RNA was used for cDNA synthesis using High-Capacity RNA-to-cDNA kit (Invitrogen). qPCR was completed with PowerUp SYBR Green Master Mix (Invitrogen) according to manufacturer protocol on a QuantStudio 6 Real Time PCR system (Applied Biosystems) and fold change was calculated using ΔΔC_t_. Primers for cDNA are listed in Supplementary Table 2 and Supplementary Table 3.

### Patient Data Gene Expression Analysis and Statistics

#### TCGA

The batch-corrected, RSEM-normalized gene level RNA-seq data from the 2018 TCGA Pan-Cancer Altas publications was used our analysis. Data was log2-transformed; and hallmark gene sets^38^ were mapped to the dataset by gene symbol and scored using ssGSEA as implemented in the GSVA R package^45^. Clinical annotations from 1100 TCGA breast samples was obtained from the cBioPortal; and used to filter the expression dataset for breast samples. Expression data was log2 transformed and median-centered; and signature genes were mapped to the dataset by gene symbol. The MYC_BC (Supplementary Table 4) and JEM^12^ MYC signatures were computed as the Pearson correlation to the directionality vector of the MYC signature genes (+1 for genes upregulated by MYC and -1 for genes downregulated by MYC); and immune cell signatures^18^ were calculated as the mean of the signature genes. Pearson correlation between the MYC_BC signature and other signatures were assessed among the 158 TNBC samples based on ER and PR status by IHC and HER status by IHC or FISH (HER2-positive if either IHC or FISH is positive).

#### ISPY-2

We computed the MYC signature score from platform-corrected, normalized, log2-scaled, median-centered pre-treatment expression data (assayed on custom Agilent 44K arrays). Signature genes were mapped to the dataset using gene symbol; and the MYC_BC signature was calculated as the Pearson correlation to the directionality vector of signature genes. A cut-point of 0 was used to dichotomize patients into High (>0) vs. Low (≤0) MYC_BC signature groups. Event-free survival was computed as time between treatment consent to loco-regional recurrence, distant recurrence or death; and patients without event were censored at time of last follow-up. Cox proportional hazard modeling was used to assess the association between the MYC signature and event-free survival (EFS) in the 28 TNBC patients from the pembrolizumab arm with available follow-up data^19^; and Kaplan Meier curves were constructed for visualization. All expression data analyses were performed using R (v 3.6.3).

### Cell Culture

MTB/TOM cells were cultured in DMEM (Gibco 11995065) supplemented with 10% FBS (Gibco) and 1% pen/strep in 5% CO_2_ at 37°C as previously described^16^. Cells were maintained in the MYC-ON state with 1 μg/mL of doxycycline (Fisher #BP2653-5) in the media, and media was changed every 2 days. MCF10A-vector (puromycin) and MCF10A-MYC cells were cultured as previously described^36^. All cells used in this study continuously tested negative for mycoplasma by PCR. For experiments with interferons, cells were seeded on day 1, treated with interferons on day 2, refreshed with new media and interferons on day 4, and harvested on day 5 (for a total of 72 hours of treatment). IFNα was used at 1,000U/mL (PBL Assay Science, #12115-1), IFNβ was used at 1,000 U/mL (PBL Assay Science, #12405-1), IFNγ was used at 100 ng/mL (Gibco, #PMC4031), and in conditions where IFNα and IFNβ were combined, 500 U/mL of each interferon was added to the well.

### Statistics for Biological Experiments

Data was analyzed in GraphPad Prism 8. Two tailed, unpaired t-test was used for comparison between treatment groups in qPCR data and immune cell flow cytometry profiling. For non-normally distributed data, such as immunohistochemistry staining and MHC-I expression, Mann-Whitney test was used. For Kaplan-Meier survival analysis, log rank test was used. Outliers were not determined, and no data were excluded in our analysis.

## Supporting information

Supplementary TablesS1-S4

## Data Availability

The MTB/TOM and MYC-driven lymphoma RNA-seq datasets were previously published^16^ and deposited to the Gene Expression Omnibus (GEO) repository (GSE130922). The RNA-seq for the MYC-driven liver model was previously published^37^ and deposited in the GEO repository GSE76078. The I-SPY2 data is previously published^19^ and available upon request with I-SPY 2 TRIAL Scientific Program Manager.

## Code Availability

The R scripts used for analyzing patient data are available by request.

## Biological Materials

The cell lines used in this study are available by request.

## Acknowledgements

This work was supported by the US National Institutes of Health 1R01CA223817 (A.G.), F32CA243548 (J.V.L), T32CA108462 (to J.V.L); the CDMRP Breast Cancer Research Program W81XWH-16-1-0603 (A.G.); the Breast Cancer Research Foundation (H.S.R.); METAvivor Research Award (A.G.); The Mark Foundation Endeavor Awards (A.G.); and the National Science Foundation #1650113 (to R.N.). We acknowledge the PFCC (RRID:SCR_018206) for assistance generating flow cytometry data. Research reported here was supported in part by the DRC Center Grant NIH P30 DK063720. We thank all of the patients who volunteered to participate in the I-SPY 2 TRIAL.

## Author Contributions

JVL, FH, MM, and AG designed the concept of the manuscript. Experiments were completed by JVL, FH, DV, RN, GH, GAH, JW, and YZ. JVL and FH completed the statistical analysis for biological experiments. CY completed and interpreted bioinformatic analysis of patient data. LJV, HSR, and LJE were involved in clinical trial design and interpreted I-SPY2 results. SS provided advocate perspectives to inform the work. JVL wrote the original manuscript, and AG, MM, DV, CY, HSR, RN, SS, and CB edited the manuscript. All authors read the manuscript and agreed to submission of the manuscript for publication.

## References

1 Doroshow, D. B. et al. PD-L1 as a biomarker of response to immune-checkpoint inhibitors. Nat Rev Clin Oncol, doi:10.1038/s41571-021-00473-5 (2021).

2 Charoentong, P. et al. Pan-cancer Immunogenomic Analyses Reveal Genotype-Immunophenotype Relationships and Predictors of Response to Checkpoint Blockade. Cell Rep 18, 248–262, doi:10.1016/j.celrep.2016.12.019 (2017).

3 Wang, S., He, Z., Wang, X., Li, H. & Liu, X. S. Antigen presentation and tumor immunogenicity in cancer immunotherapy response prediction. Elife 8, doi:10.7554/eLife.49020 (2019).

4 Samstein, R. M. et al. Tumor mutational load predicts survival after immunotherapy across multiple cancer types. Nat Genet 51, 202–206, doi:10.1038/s41588-018-0312-8 (2019).

5 Yarchoan, M., Hopkins, A. & Jaffee, E. M. Tumor Mutational Burden and Response Rate to PD-1 Inhibition. N Engl J Med 377, 2500–2501, doi:10.1056/NEJMc1713444 (2017).

6 Li, J. et al. Epigenetic driver mutations in ARID1A shape cancer immune phenotype and immunotherapy. J Clin Invest, doi:10.1172/JCI134402 (2020).

7 Loi, S. et al. RAS/MAPK Activation Is Associated with Reduced Tumor-Infiltrating Lymphocytes in Triple-Negative Breast Cancer: Therapeutic Cooperation Between MEK and PD-1/PD-L1 Immune Checkpoint Inhibitors. Clin Cancer Res 22, 1499–1509, doi:10.1158/1078-0432.CCR-15-1125 (2016).

8 Haikala, H. M. et al. Pharmacological reactivation of MYC-dependent apoptosis induces susceptibility to anti-PD-1 immunotherapy. Nat Commun 10, 620, doi:10.1038/s41467-019-08541-2 (2019).

9 Han, H. et al. Small-Molecule MYC Inhibitors Suppress Tumor Growth and Enhance Immunotherapy. Cancer Cell 36, 483–497 e415, doi:10.1016/j.ccell.2019.10.001 (2019).

10 Kortlever, R. M. et al. Myc Cooperates with Ras by Programming Inflammation and Immune Suppression. Cell 171, 1301–1315 e1314, doi:10.1016/j.cell.2017.11.013 (2017).

11 Cancer Genome Atlas, N. Comprehensive molecular portraits of human breast tumours. Nature 490, 61–70, doi:10.1038/nature11412 (2012).

12 Horiuchi, D. et al. MYC pathway activation in triple-negative breast cancer is synthetic lethal with CDK inhibition. J Exp Med 209, 679–696, doi:10.1084/jem.20111512 (2012).

13 D’Cruz, C. M. et al. c-MYC induces mammary tumorigenesis by means of a preferred pathway involving spontaneous Kras2 mutations. Nat Med 7, 235–239, doi:10.1038/84691 (2001).

14 Casey, S. C. et al. MYC regulates the antitumor immune response through CD47 and PD-L1. Science 352, 227–231, doi:10.1126/science.aac9935 (2016).

15 Pfefferle, A. D. et al. Transcriptomic classification of genetically engineered mouse models of breast cancer identifies human subtype counterparts. Genome Biol 14, R125, doi:10.1186/gb-2013-14-11-r125 (2013).

16 Rohrberg, J. et al. MYC Dysregulates Mitosis, Revealing Cancer Vulnerabilities. Cell Rep 30, 3368–3382 e3367, doi:10.1016/j.celrep.2020.02.041 (2020).

17 Chandriani, S. et al. A core MYC gene expression signature is prominent in basal-like breast cancer but only partially overlaps the core serum response. PLoS One 4, e6693, doi:10.1371/journal.pone.0006693 (2009).

18 Danaher, P. et al. Gene expression markers of Tumor Infiltrating Leukocytes. J Immunother Cancer 5, 18, doi:10.1186/s40425-017-0215-8 (2017).

19 Nanda, R. et al. Effect of Pembrolizumab Plus Neoadjuvant Chemotherapy on Pathologic Complete Response in Women With Early-Stage Breast Cancer: An Analysis of the Ongoing Phase 2 Adaptively Randomized I-SPY2 Trial. JAMA Oncol 6, 676–684, doi:10.1001/jamaoncol.2019.6650 (2020).

20 Fu, J. et al. Large-scale public data reuse to model immunotherapy response and resistance. Genome Med 12, 21, doi:10.1186/s13073-020-0721-z (2020).

21 Jiang, P. et al. Signatures of T cell dysfunction and exclusion predict cancer immunotherapy response. Nat Med 24, 1550–1558, doi:10.1038/s41591-018-0136-1 (2018).

22 Mariathasan, S. et al. TGFbeta attenuates tumour response to PD-L1 blockade by contributing to exclusion of T cells. Nature 554, 544–548, doi:10.1038/nature25501 (2018).

23 Miao, D. et al. Genomic correlates of response to immune checkpoint therapies in clear cell renal cell carcinoma. Science 359, 801–806, doi:10.1126/science.aan5951 (2018).

24 Schaub, F. X. et al. Pan-cancer Alterations of the MYC Oncogene and Its Proximal Network across the Cancer Genome Atlas. Cell Syst 6, 282–300 e282, doi:10.1016/j.cels.2018.03.003 (2018).

25 Robertson, A. G. et al. Comprehensive Molecular Characterization of Muscle-Invasive Bladder Cancer. Cell 171, 540–556 e525, doi:10.1016/j.cell.2017.09.007 (2017).

26 Bailey, S. T. et al. MYC activation cooperates with Vhl and Ink4a/Arf loss to induce clear cell renal cell carcinoma. Nat Commun 8, 15770, doi:10.1038/ncomms15770 (2017).

27 Cornel, A. M., Mimpen, I. L. & Nierkens, S. MHC Class I Downregulation in Cancer: Underlying Mechanisms and Potential Targets for Cancer Immunotherapy. Cancers (Basel) 12, doi:10.3390/cancers12071760 (2020).

28 Zaretsky, J. M. et al. Mutations Associated with Acquired Resistance to PD-1 Blockade in Melanoma. N Engl J Med 375, 819–829, doi:10.1056/NEJMoa1604958 (2016).

29 McGranahan, N. et al. Allele-Specific HLA Loss and Immune Escape in Lung Cancer Evolution. Cell 171, 1259–1271 e1211, doi:10.1016/j.cell.2017.10.001 (2017).

30 Gettinger, S. et al. Impaired HLA Class I Antigen Processing and Presentation as a Mechanism of Acquired Resistance to Immune Checkpoint Inhibitors in Lung Cancer. Cancer Discov 7, 1420–1435, doi:10.1158/2159-8290.CD-17-0593 (2017).

31 Torrejon, D. Y. et al. Overcoming Genetically Based Resistance Mechanisms to PD-1 Blockade. Cancer Discov 10, 1140–1157, doi:10.1158/2159-8290.CD-19-1409 (2020).

32 Versteeg, R., Noordermeer, I. A., Kruse-Wolters, M., Ruiter, D. J. & Schrier, P. I. c-myc down-regulates class I HLA expression in human melanomas. EMBO J 7, 1023–1029 (1988).

33 Bernards, R., Dessain, S. K. & Weinberg, R. A. N-myc amplification causes down-modulation of MHC class I antigen expression in neuroblastoma. Cell 47, 667–674, doi:10.1016/0092-8674(86)90509-x (1986).

34 Ludigs, K. et al. NLRC5 exclusively transactivates MHC class I and related genes through a distinctive SXY module. PLoS Genet 11, e1005088, doi:10.1371/journal.pgen.1005088 (2015).

35 Meissner, T. B. et al. NLR family member NLRC5 is a transcriptional regulator of MHC class I genes. Proc Natl Acad Sci U S A 107, 13794–13799, doi:10.1073/pnas.1008684107 (2010).

36 Martins, M. M. et al. Linking tumor mutations to drug responses via a quantitative chemical-genetic interaction map. Cancer Discov 5, 154–167, doi:10.1158/2159-8290.CD-14-0552 (2015).

37 Kress, T. R. et al. Identification of MYC-Dependent Transcriptional Programs in Oncogene-Addicted Liver Tumors. Cancer Res 76, 3463–3472, doi:10.1158/0008-5472.CAN-16-0316 (2016).

38 Liberzon, A. et al. The Molecular Signatures Database (MSigDB) hallmark gene set collection. Cell Syst 1, 417–425, doi:10.1016/j.cels.2015.12.004 (2015).

39 Nistico, P. et al. Effect of recombinant human leukocyte, fibroblast, and immune interferons on expression of class I and II major histocompatibility complex and invariant chain in early passage human melanoma cells. Cancer Res 50, 7422–7429 (1990).

40 Klinman, D. M. Immunotherapeutic uses of CpG oligodeoxynucleotides. Nat Rev Immunol 4, 249–258, doi:10.1038/nri1329 (2004).

41 Sagiv-Barfi, I. et al. Eradication of spontaneous malignancy by local immunotherapy. Sci Transl Med 10, doi:10.1126/scitranslmed.aan4488 (2018).

42 Muthalagu, N. et al. Repression of the Type I Interferon pathway underlies MYC & KRAS-dependent evasion of NK & B cells in Pancreatic Ductal Adenocarcinoma. Cancer Discov, doi:10.1158/2159-8290.CD-19-0620 (2020).

43 Burr, M. L. et al. An Evolutionarily Conserved Function of Polycomb Silences the MHC Class I Antigen Presentation Pathway and Enables Immune Evasion in Cancer. Cancer Cell 36, 385–401 e388, doi:10.1016/j.ccell.2019.08.008 (2019).

44 Topper, M. J. et al. Epigenetic Therapy Ties MYC Depletion to Reversing Immune Evasion and Treating Lung Cancer. Cell 171, 1284–1300 e1221, doi:10.1016/j.cell.2017.10.022 (2017).

45 Hanzelmann, S., Castelo, R. & Guinney, J. GSVA: gene set variation analysis for microarray and RNA-seq data. BMC Bioinformatics 14, 7, doi:10.1186/1471-2105-14-7 (2013).

