## Supplementary_Figures for "Combinatorial Immunotherapies Overcome MYC-Driven Immune Evasion"

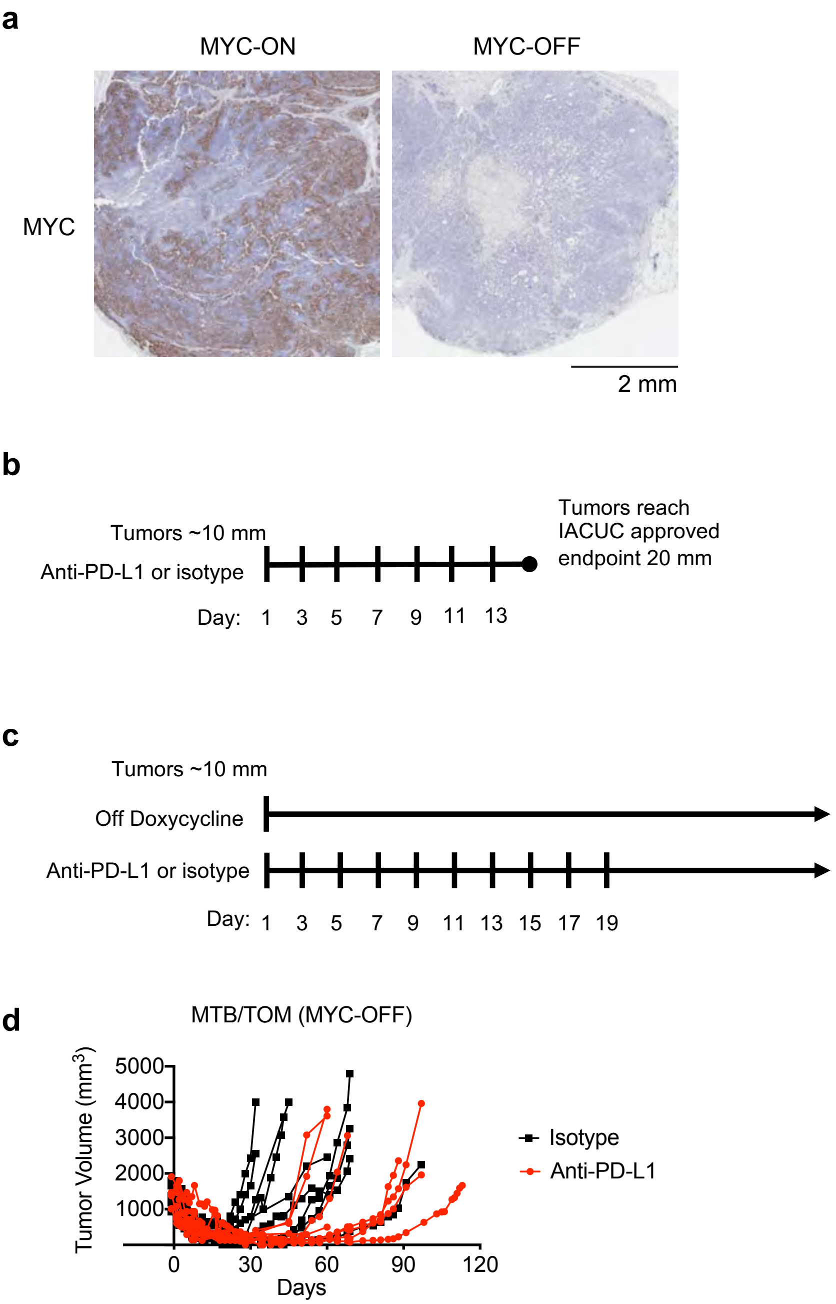


**Extended Data Fig. 1**: **The MYC-driven TNBC model for testing anti-PD-L1. (a)** MTB/TOM tumors downregulate MYC upon doxycycline withdrawal**.** Immunohistochemistry staining for MYC in tumors from an animal on doxycycline chow (MYC-ON) or an animal 3 days off doxycycline chow (MYC-OFF). **(b)** Animals were randomized into treatment arms once tumors reached approximately 10 mm in length (longest length in any direction). Anti-PD-L1 or isotype control antibody was administered on days marked by the tick marks. **(c)** Animals were taken off doxycycline and enrolled for anti-PD-L1 or isotype. Animals remained off doxycycline for the duration of the study. **(d)** Tumor volume starting on day 1, the day doxycycline chow is removed (MYC-OFF state) and anti-PD-L1 therapy (n=8) or isotype control antibody (n=9) is initiated.

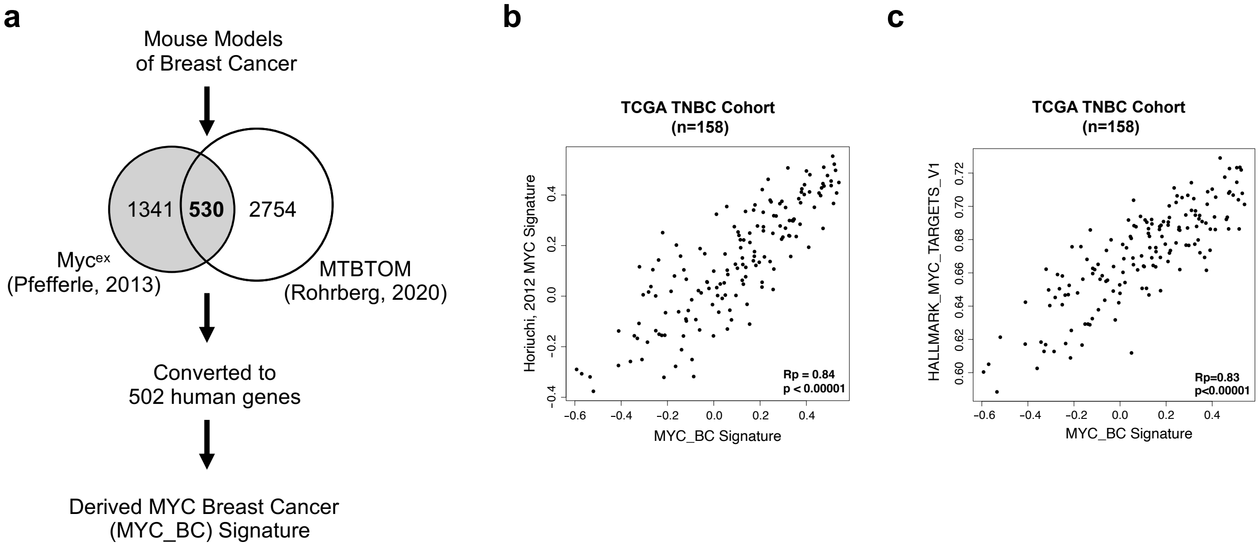


**Extended Data Fig. 2: *In vivo* derived MYC signature (MYC_BC) correlates with published *in vitro* derived MYC signatures. (a)** Diagram depicting derivation of the MYC Breast Cancer (MYC_BC) Signature. **(b)** Scatterplot correlation between MYC_BC signature expression and 352 gene MYC signature (Horiuchi, 2012; Chandriani, 2009). **(c)** Scatterplot correlation between MYC_BC signature expression and MSigDB MYC v1 gene set signature. TCGA TNBC cohort (n=158). Rp, Pearson's correlation coefficient

### TCGA TNBC (n=158)

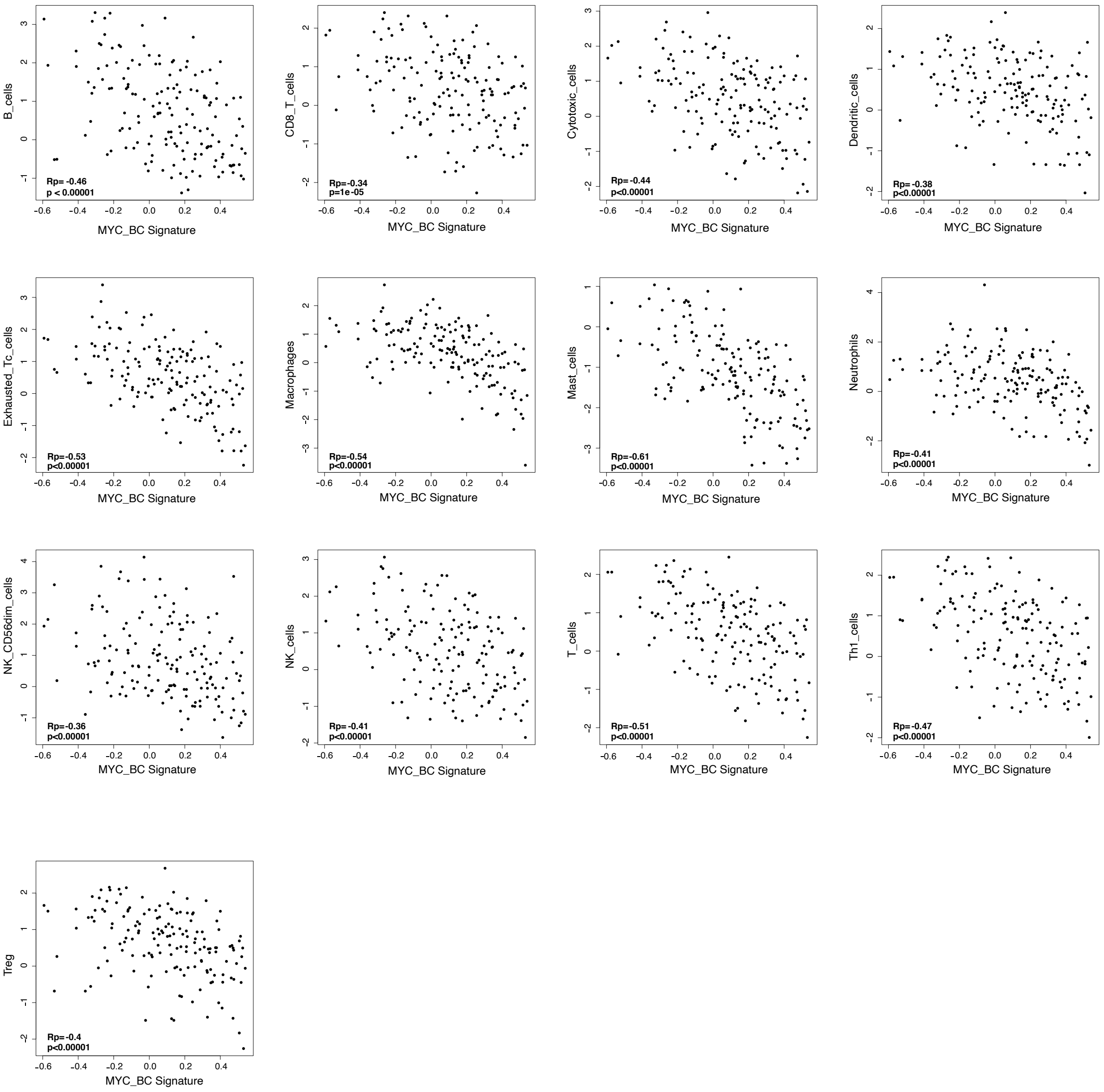


**Extended Data Fig. 3: MYC_BC signature is anti-correlated with multiple immune cell signatures.** Scatterplot correlation between MYC_BC signature expression and published immune signatures from Danaher, 2017. TCGA TNBC cohort (n=158). Rp, Pearson's correlation coefficient.

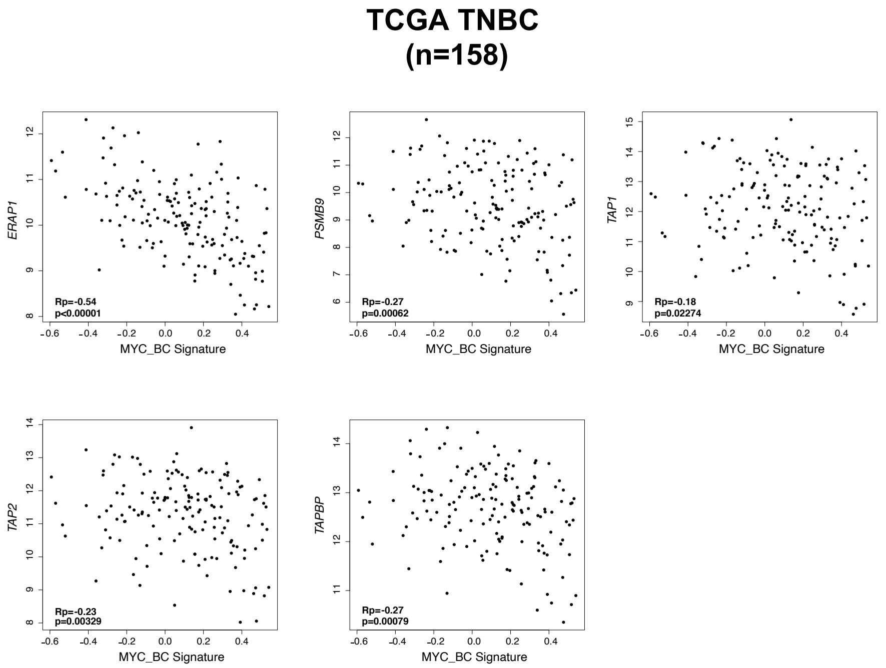


**Extended Data Fig. 4: MYC_BC signature is anti-correlated with genes required for MHC-I antigen presentation.** Scatterplot correlation between MYC_BC signature expression and expression score for each gene in TCGA TNBC cohort (n=158). Rp, Pearson's correlation coefficient.

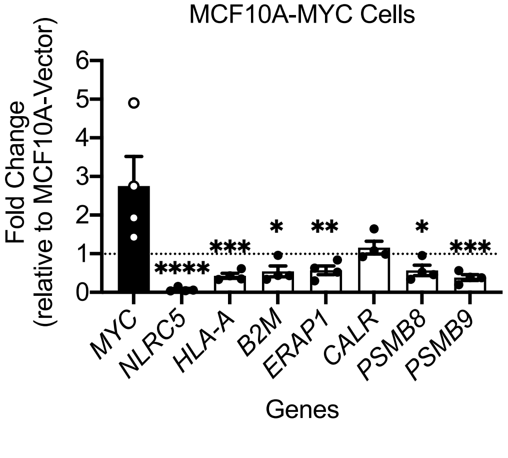


**Extended Data Fig. 5: MYC overexpression leads to lower expression of antigen presentation genes.** Multiple genes are significantly changed in MCF10A-MYC cells. Each gene is relative to expression vector control MCF10A cells, with fold change less than 1 (dotted line) demonstrating gene repression. Shown here are n=4 independent cell passage replicates. Mean± S.E.M, unpaired t-test. ∗, $p < 0.05$; ∗∗, $p < 0.01$; ∗∗∗, $p < 0.001$; ∗∗∗∗, $p < 0.0001$.

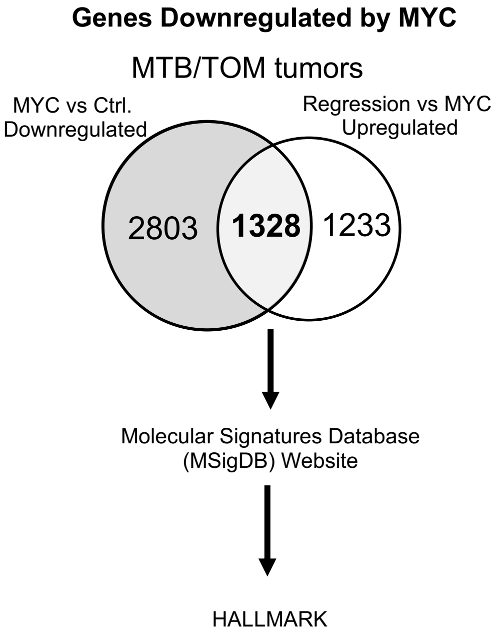


**Extended Data Fig. 6: Identification of genes downregulated by MYC.** The differential expression data was downloaded from Rohrberg, 2020 (GSE130922). Genes with a 2-fold change in expression and an adjusted p-value < 0.05 were selected from the published analysis. The two lists were compared using online tool Venny 2.0 (https://bioinfogp.cnb.csic.es/tools/venny/index.html) Overlapping genes were entered into GSEA/MSigDB (https://www.gsea-msigdb.org/gsea/msigdb) for evaluation of significant overlap with top 10 MSigDB HALLMARK collection.

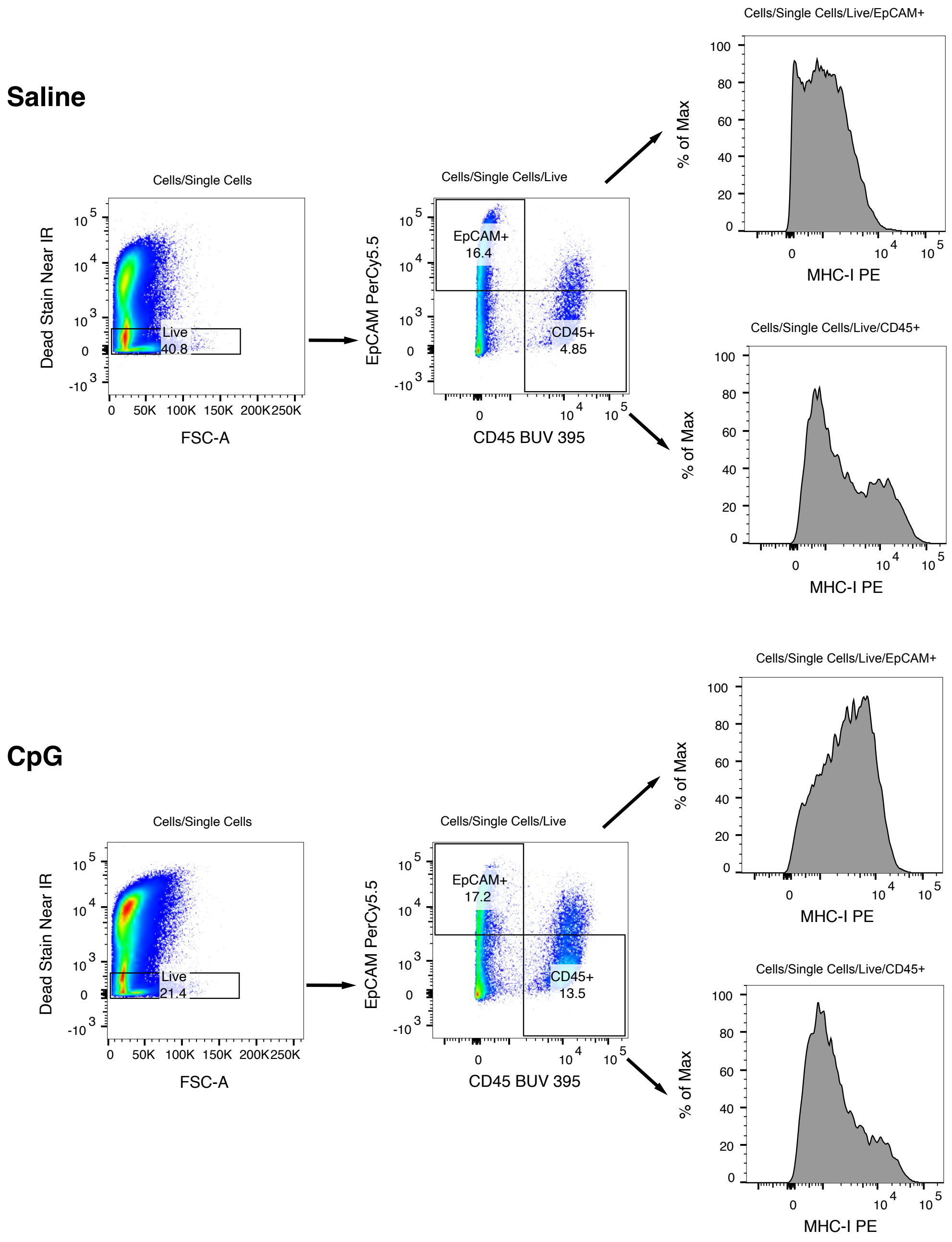


**Extended Data Fig. 7: Gating strategy for MHC-I in CpG treated tumors**. Cells were gated on size and granularity (SSC-A, FSC-A), Singlets (FSC-H, FSC-A), Live (negative stain), and EPCAM+, CD45- or EPCAM-, CD45+.

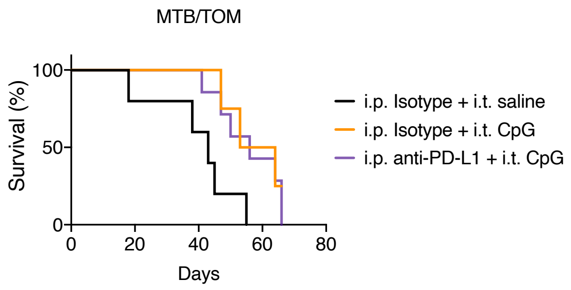


**Extended Data Fig. 8: CpG+anti-PD-L1 does not extend survival beyond anti-PD-L1 alone.** Survival analysis for MTB/TOM treated intratumoral (i.t.) saline or CpG and intraperitoneal (i.p.) anti-PD-L1 or isotype control antibody. Median survival was 43 days, 58.5 days and 56 days for Isotype + saline (n=5), isotype + CpG (n=4), and anti-PD-L1+CpG (n=7), respectively.

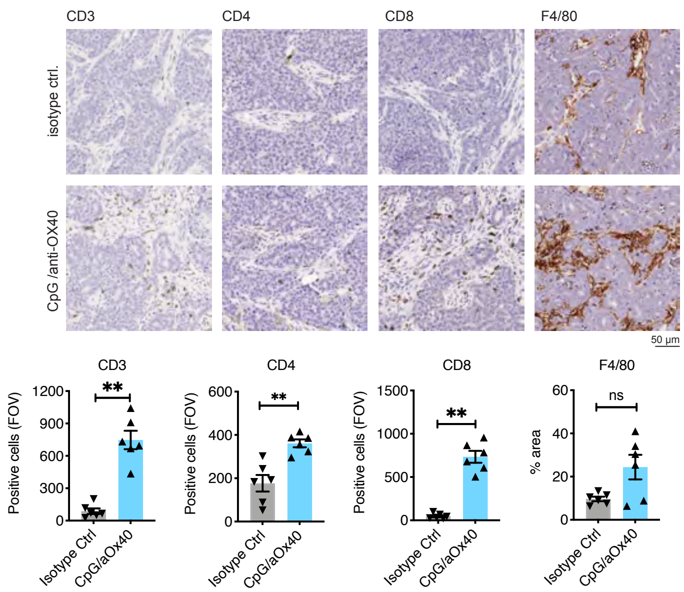


**Extended Data Fig. 9: CpG/aOX40 increases T-cell infiltration.** Immunohistochemistry staining for immune cells in the tumor on day 10 (top) and bar graph (below) depicting counts per field of view (FOV). Images taken at 40X. n=2 animals per group, 3 FOV analyzed per tumor, mean ± S.E.M., Mann-Whitney test. ✱✱, $p < 0.01$

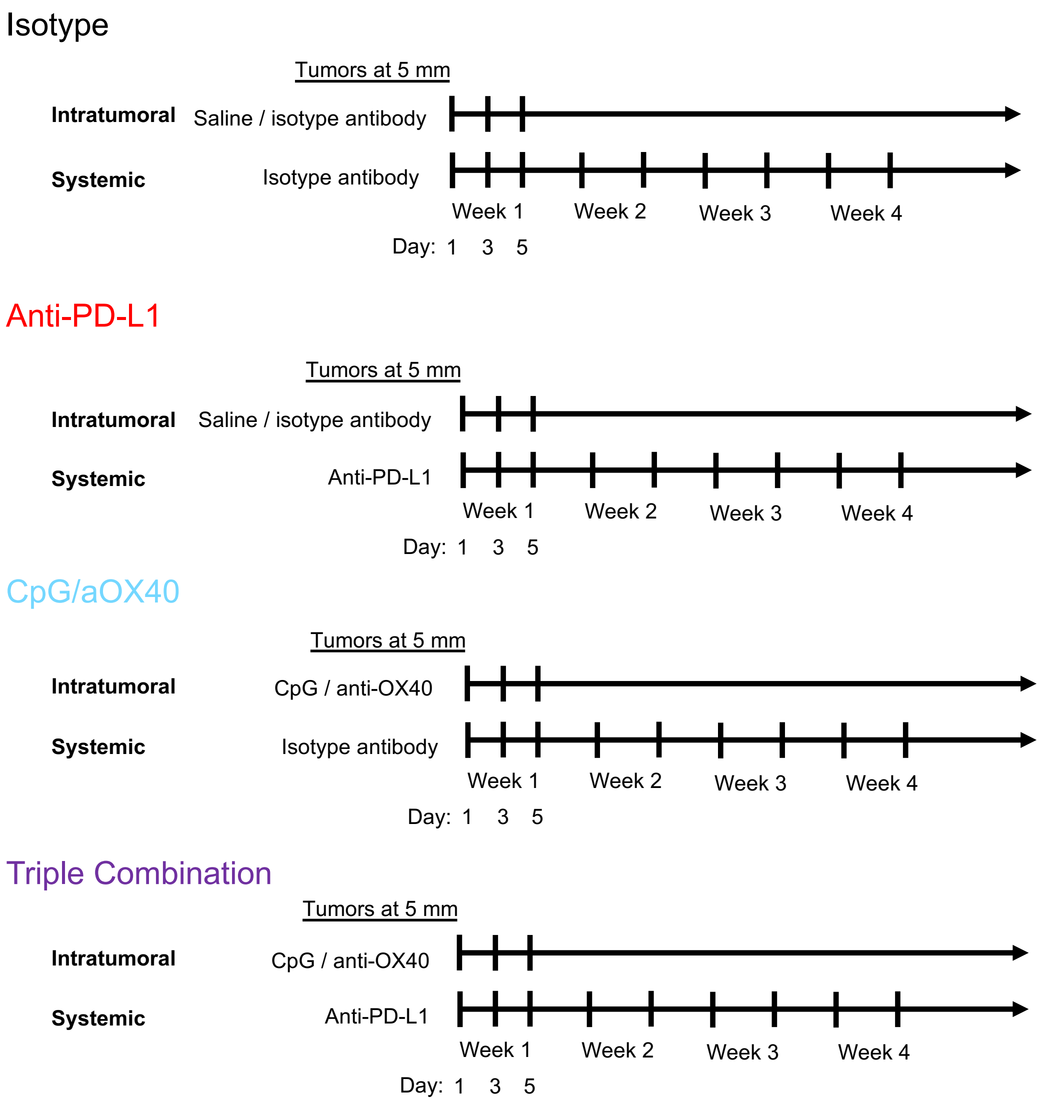


**Extended Data Fig. 10: Treatment schedule related to Figure 3h, i.** Animals were randomized into treatment arms once tumors reached 5 mm in length (longest length in any direction). All agents were administered on days 1, 3, and 5, followed by maintenance on anti-PD-L1 or isotype injections twice a week for 6 additional doses (total 9 doses).

Extended Data Table 1: Pathways enriched in MYC-downregulated genes

| **10 Most Downregulated Hallmark Signatures** | | |
|---|---|---|
| Gene Set Name | k/K | FDR q-value |
| HALLMARK_INTERFERON_GAMMA_RESPONSE | 0.32 | 2.95E-46 |
| HALLMARK_INTERFERON_ALPHA_RESPONSE | 0.351 | 3.50E-26 |
| HALLMARK_ALLOGRAFT_REJECTION | 0.215 | 1.37E-23 |
| HALLMARK_COMPLEMENT | 0.165 | 1.22E-14 |
| HALLMARK_INFLAMMATORY_RESPONSE | 0.16 | 6.51E-14 |
| HALLMARK_APOPTOSIS | 0.168 | 2.17E-12 |
| HALLMARK_TNFA_SIGNALING_VIA_NFKB | 0.145 | 1.11E-11 |
| HALLMARK_APICAL_JUNCTION | 0.14 | 4.96E-11 |
| HALLMARK_EPITHELIAL_MESENCHYMAL_TRANSITION | 0.14 | 4.96E-11 |
| HALLMARK_IL2_STAT5_SIGNALING | 0.135 | 2.05E-10 |
